# Assessment of endemic northern swamp deer (*Rucervus duvaucelii duvaucelii*) distribution and identification of priority conservation areas through modeling and field surveys across north India

**DOI:** 10.1101/2020.07.09.194803

**Authors:** Shrutarshi Paul, Debanjan Sarkar, Abhilash Patil, Tista Ghosh, Gautam Talukdar, Mukesh Kumar, Bilal Habib, Parag Nigam, Dhananjai Mohan, Bivash Pandav, Samrat Mondol

## Abstract

Recent declines in large herbivores have led to significant conservation efforts globally. However, the niche-specific megaherbivores residing outside protected areas face more imminent extinction threats. Swamp deer, the obligate grassland-dwelling endemic cervid is the most extinction-prone megaherbivore in the Indian subcontinent. Limited information on distribution and habitat status pose significant conservation and management challenges for the remaining fragmented populations in north, north-east and central India. To this end, we combined exhaustive field surveys and Maximum Entropy (MaxEnt) modeling to generate the most detailed distribution map for the northern swamp deer subspecies. We used primary data from more than 6000 km^2^ field surveys and eight ecologically relevant covariates for model predictions. Grassland cover, annual mean temperature and distance from water were the major factors that predicted the species distribution. Models predicted swamp deer distribution in only ~3% of the entire landscape, covering both protected (~1.4%) as well as non-protected (~1.6%) areas. Our validation surveys in some of these predicted areas confirmed swamp deer presence and indicated ~85% model accuracy. Finally, we identified four ‘Priority Conservation Areas’ still retaining adequate grassland habitat and species presence that require immediate attention to ensure population connectivity across this landscape. These results highlight the importance of the marginalized grassland ecosystems of northern India that still retains high biodiversity. We suggest a swamp deer-centric conservation approach to protect these human-dominated habitats and emphasize in generating such information for other endemic, habitat-specialist species across the globe.

## Introduction

Large herbivores play critical roles in shaping ecosystems through various ecological functions (nutrient cycling, seed dispersal etc.) that affect forest structure, regeneration and benefit other species (Danell et al., 2006; Ripple et al., 2015). In recent time, habitat destruction, resource depletion, hunting and human-animal conflict have resulted in massive declines of the world’s large herbivores (Ceballos et al., 2017; Lindsey et al., 2017; Ripple et al., 2015) such that about 60% of them are threatened with extinction (Ripple et al., 2015; Trouwborst, 2019). The effects of such anthropogenic factors towards their extinction are further exacerbated by their relatively low population densities, unique habitat requirements and slow life history characteristics (Ripple et al., 2016; Wallach et al., 2015). While significant global conservation efforts helped in population recovery of some of the charismatic large herbivores (African rhinoceros-Emsile and Brooks, 1999; Asian elephant, gaur and sambar- Madhusudan, 2004; black rhinoceros- Brodie et al., 2011; Asian elephant and one-horned rhinoceros- Williams, 2011; elk, European bison and red deer- Sylven et al., 2012), it has mostly been targeted within protected areas, that represents only ~13% of the total land surface area (Chape et al., 2005; UNEP-WCMC, 2012). The challenges are even more acute for habitat-specialist large herbivores that are distributed outside protected areas along the human-livestock-wildlife interfaces, where implementing regular management efforts like the protected areas are difficult (Karanth et al., 2010; Kshettry et al., 2020). Although these non-protected areas harbour significant mammalian biodiversity (Dorji et al., 2019; Harihar, 2011; Punjabi and Rao, 2017) and act as important corridor habitats for free-ranging animals (Neelakantan et al., 2019; Sunderraj et al., 1995), they are often overexploited (land conversion, grazing, unregulated poaching and hunting practices etc.) (Macdonald et al., 2013; Ripple et al., 2015, 2014) and experience strong human-wildlife conflict (crop raiding, damage to infrastructures and injuries to humans) (Gubbi et al., 2014; Kissui, 2008). Conservation of such large herbivores in these non-protected areas is dependent on accurate information on distribution, habitat status and other biological parameters.

The Indian subcontinent retains one of the highest diversity of large herbivores in south and south-east Asia (Johnsingh et al., 2004; Ahrestani et al., 2011). The large herbivore species assemblages are currently facing serious conservation challenges in the subcontinent with 12 of the 15 terrestrial large herbivores classified as ‘‘Threatened’’ by IUCN (Ripple et al., 2014, 2015). A recent study by Karanth et al. (2010) has shown that within the subcontinent, the habitat-specific large herbivores (for example, swamp deer, rhino, wild buffalo etc.) are much more prone to extinction due to their inherent species biology and various anthropogenic activities and require accurate information on distribution, population size and various habitat parameters to ensure their future survival. However, generating such information is challenging for many of these species due to their low density and cryptic behaviour, particularly for those living in a mosaic of protected and non-protected areas surrounding human settlements (Jathanna et al., 2003; Linkie et al., 2013; Marshal, 2016). Given that only about 5% of land comes under protected area category within India (Wildlife Institute of India, 2020), it is critical to generate detailed information on distribution of these habitat-specialist large herbivores and identify important habitats outside protected areas for their long-term conservation.

The swamp deer or barasingha (*Rucervus duvaucelli*) exemplify such habitat-specialist endemic large herbivores of the south Asian region (Qureshi et al., 2004; Tewari and Rawat, 2013). This obligate swampy grassland-dwelling cervid was historically distributed throughout the Indo-Gangetic plains and the lowlands spanning across the southern Himalayas covering Pakistan, southern Nepal, India and Bangladesh (Groves, 1982; Sankaran, 1989; Schaller, 1967). However, it is now classified as “Vulnerable” according to IUCN due to severe population decline and range contraction during last century (Duckworth et al., 2015). This has resulted in a global population size of <5000 individuals, restricted to isolated populations in north, north-east and central India and south-west Nepal (Qureshi et al., 2004; Tewari and Rawat, 2013). Among the three known subspecies (northern-*Rucervus duvaucelii duvaucelii*, central-*Rucervus duvaucelii branderi* and eastern-*Rucervus duvaucelii ranjitsinhi*), the northern subspecies retains about 80% of the global population (Qureshi et al., 2004, 1995) and is distributed as small, fragmented populations across the states of Uttar Pradesh, Uttarakhand in India and southwestern Nepal (Khan and Khan, 1999; Paul et al., 2018). The Indian part of the northern swamp deer distribution is broadly divided into two habitat blocks: a) the habitat block along the Sharda river basin in the state of Uttar Pradesh covering a mosaic of protected (Pilibhit Tiger Reserve, Kishanpur Wildlife Sanctuary, Dudhwa National Park and Katerniaghat Wildlife Sanctuary) and unprotected areas surrounding them and b) habitat block along river Ganga covering protected areas in Uttarakhand (Jhilmil Jheel Conservation Reserve) and Uttar Pradesh (Hastinapur Wildlife Sanctuary) along with surrounding unprotected, patchy grassland habitats (Qureshi et al., 2004, 1995) (Figure 1). Encroachment pressures on the grasslands (only 2% of the entire Gangetic floodplains) (Dinerstein, 2003) from one of the most highly dense human populations (about 800 people/km^2^ compared to 382 people/km^2^ in India, Census of India 2011) living in this region and scanty information on exact distribution and habitat of swamp deer status pose significant conservation challenges (Duckworth, et al., 2015; Midha and Mathur, 2010; Paul et al., 2018). A recent study by Paul et al. (2018) has reported a detailed assessment of the subspecies presence and habitat status in the Gangetic block, with previously unreported swamp deer harbouring areas. This indicates promising signs for this subspecies, but a landscape wide assessment of swamp deer distribution is essential for developing better management plans for the species and its remaining grassland habitats.

**Figure 1:**
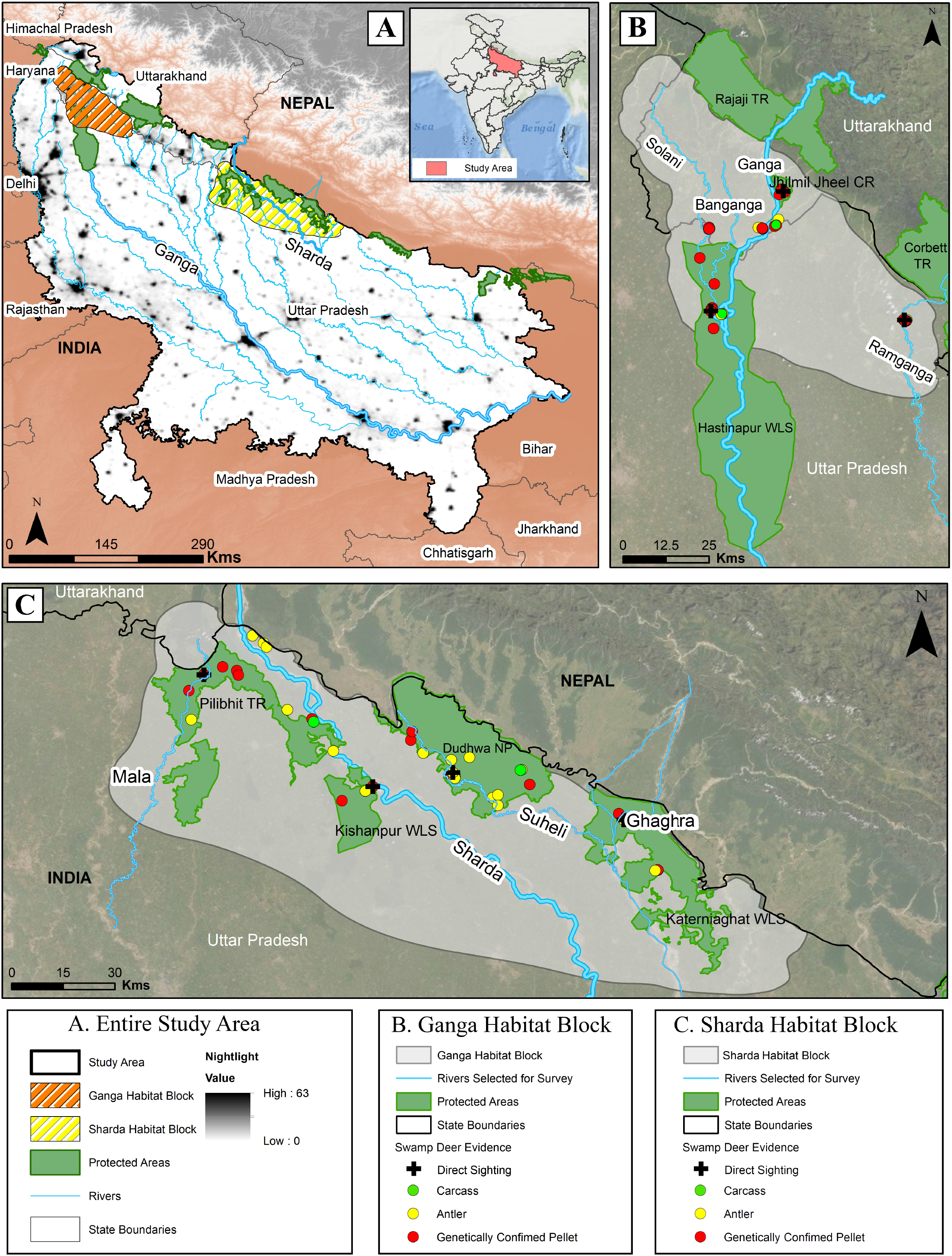
The study area map across the states of Uttarakhand (selected area) and Uttar Pradesh, north India. The map shows (A) Entire study area covering both Ganga and Sharda Habitat Blocks; (B) the locations of the confirmed swamp deer evidences within Ganga Habitat Block (data from Paul et al. 2018) and (C) the locations of the confirmed swamp deer evidences within Sharda Habitat Block. Different evidence types (direct and indirect) are presented in the figure legends.

However, conducting surveys in this human-dominated landscape is logistically challenging and probably requires a combination of different approaches to assess the fine-scale distribution pattern of this subspecies.

In this study, we assessed *Rucervus duvaucelii duvaucelii* distribution covering protected and non-protected areas across the upper Gangetic plains, within the boundaries of Uttarakhand and Uttar Pradesh. We used multidisciplinary approaches involving scientifically designed rigorous field surveys in Ganga (data from Paul et al., 2018) and Sharda block, distribution modeling and subsequent field validations to assess current swamp deer distribution and status of their habitats. Further, we have identified four new conservation priority areas that are crucial for future swamp deer conservation planning. The findings of this study would be important in developing long-term conservation strategies for swamp deer and the wetland/grassland habitats that hosts a number of other threatened fauna.

## Methods

### Research permissions

All required permissions for swamp deer surveys and biological sample collection were provided by the Forest Departments of Uttarakhand and Uttar Pradesh (Letter no: 90/5, 978/6-32/56, 1127/23-2-12(G) and 2233/23-2-12(G)). Due to the non-invasive nature of sampling, no ethical clearance was required in this study.

### Study Area

This study was conducted covering about 253188 km^2^ area across selected districts of Uttarakhand (Dehradun, Haridwar, Pauri-Garhwal, Nainital and Udham Singh Nagar) and the entire state of Uttar Pradesh (Figure 1). The area covers the floodplains of the Ganga river and a number of its tributaries like Sharda, Ghaghra, Gandak, Khoh, Ramganga etc. (Figure 1) including a complex network of riverine-grassland and grassland-forest systems. This region hosts the only existing populations of northern swamp deer (*Rucervus duvaucelii duvaucelii*) in India (Qureshi et al., 2004). The primary vegetation in the grassland patches comprises *Typha* sp., *Phragmites* sp. and *Saccharum* spp. (Tewari and Rawat, 2013). Larger animals in the study area include swamp deer (*Rucervus duvaucelii*), hog deer (*Axis porcinus*), spotted deer (*Axis axis*), nilgai (*Boselaphus tragocamelus*), smooth-coated otter (*Lutrogale perspicillata*), fishing cat (*Prionailurus viverrinus*) and iconic wetland birds such as sarus crane (*Antigone antigone*), black-necked stork (*Ephippiorhynchus asiaticus*), lesser adjutant (*Leptoptilos javanicus*), Pallas’s fish eagle (*Haliaeetus leucoryphus*) and bar-headed goose (*Anser indicus*) (Grimmett et al., 2013). The area experiences extreme seasonal variations in temperature (7°C-40°C) and receives an average annual rainfall of 948 mm (502.4-2113.9 mm) with occasional flooding during July-September (Verma et al., 2018).

### Species Distribution Modeling (SDM) for swamp deer using MaxEnt

#### Generation of swamp deer occurrence points

We used MaxEnt (Maximum Entropy) species distribution algorithm version 3.3.3k (Phillips et al., 2006) to model potential distribution of swamp deer in the study area. Maximum Entropy is one of the most widely used algorithms for species distribution with presence-only data (Elith et al., 2006). For best performance of the algorithm, it is critical to have unbiased, error-free species presence points for initial calibration of the models (Kramer-Schadt et al., 2013). Biased survey design including inadequate survey efforts, sporadic or clumped location information, erroneous presence data etc. affect model accuracy (Costa et al., 2015; Kramer-Schadt et al., 2013). We conducted systematic intensive surveys to generate accurate swamp deer presence points from both Ganga and Sharda river habitat blocks for model calibration. Details of the survey and fine-scale swamp deer distribution at the Gangetic habitat block from Jhilmil Jheel Conservation Reserve (JJCR) in Uttarakhand to Bijnor Barrage area of Hastinapur Wildlife Sanctuary (HWLS) in Uttar Pradesh is described in Paul et al. (2018). In Sharda block, swamp deer presence has been reported from a few well-known swamps within protected areas only. For a comprehensive detailed assessment of swamp deer distribution in Sharda block, we adopted a similar combined approach of opportunistic unstructured questionnaire surveys (see details in Paul et al., 2018) and Google Earth imageries (Hu et al., 2013) to locate possible areas (both protected and adjoining non-protected) where the species was found. As most of the potential habitats in this region are protected, the questionnaire surveys were targeted towards Forest Department staff and occasionally with local cattle herders, boatmen and fishermen. All information indicated that majority of the grassland patches are currently restricted within 10 km areas on both banks of the rivers Mala (Pilibhit Tiger Reserve), Sharda (Pilibhit Tiger Reserve and Kishanpur Wildlife Sanctuary), Suheli (Dudhwa National Park) and Ghaghra (Katerniaghat Wildlife Sanctuary). Subsequently, we intensively surveyed both banks of the entire stretch of these rivers covering both protected as well as non-protected areas. The surveys were conducted using vehicle, boat or on foot depending on the area during the months of May of 2016 and 2017 covering a total area of around 6000 km^2^.

During surveys, we collected geo-tagged information on direct (sighting, carcass etc.) as well as indirect (antler and genetically identified dung pellets) evidence of swamp deer presence. Swamp deer antlers are quite distinctive from other coexisting cervids in this landscape (sambar, hog deer, chital and barking deer) (Paul et al., 2019). However, swamp deer pellets are difficult to identify based on morphology and we conducted molecular species identification using species-specific primers for different cervids described in Paul et al. (2019). Remaining non-amplified samples were tested with ungulate specific primers (Gupta et al., 2014) to identify species.

#### Fine-tuning of Species occurrence data for model calibration

Combining the earlier records from upper Gangetic habitat block (n=169, Paul et al., 2018) and new surveys at Sharda basin habitat block (n=139, 2 carcasses, 7 direct sightings, 51 genetically identified pellets and 79 antlers, Supplementary Table 2), we gathered 308 confirmed swamp deer location points from the entire landscape. Upon plotting these locations in a map, we found that many confirmed swamp deer locations were clustered within the fragmented grassland patches, leading to spatial autocorrelation (around 4.4 points/ km^2^ grid). Such spatial patterns may lead to overfit model response to covariates, thereby limiting the model’s prediction accuracy (Boria et al., 2014; Kramer-Schadt et al., 2013). Calibration and evaluation localities that are close to each other might lead to inflated values of performance (Veloz, 2009; Hijmans, 2012). To reduce such effects, we selected a single presence record in every one km^2^ area (as the species is mostly restricted to habitat patches of around one km^2^ currently within the grassland-cropland mosaics within this landscape) using Spatially Rarefy Tool of SDM tool Box (Brown, 2014) in ArcGIS 10.2.2. We used 69 such records for species distribution modeling from a total of 308 points in the Gangetic plains of Uttarakhand and Uttar Pradesh.

#### Selection of covariates

Based on swamp deer species biology, we initially selected numerous climatic, vegetation,topographic and anthropogenic variables to model swamp deer distribution. Bioclimatic variables are regularly used in defining species environmental niches (Booth et al., 2014; Bradie and Leung, 2017) and we downloaded 30 arc-second spatial resolution (or one km^2^ spatial resolution) data of four such variables from WorldClim-Global Climate Data (www.worldclim.org/bioclim). The topographic covariates in the form of “Elevation” and “Slope” layers were downloaded from the USGS website (www.usgs.gov). The “slope” was calculated using ArcToolbox, Spatial Analyst in ArcGIS 10.2 (ESRI, Redlands, CA, USA). We also selected Normalized Difference Vegetation Index (NDVI) of March and May 2017, distance from water, nightlight and human population density (anthropogenic variables), rescaled to one km^2^ resolution as covariates (Table 1).

**Table 1:** Details of all covariates selected for modelling distribution of swamp deer

| Category | Variables | Data source | Abbreviation | Units/Range | Variables selected |
| --- | --- | --- | --- | --- | --- |
| Bioclimatic | Annual Mean Temperature | WorldClim (https://worldclim.org/data/bioclim.html) | Bio1 | °C | Yes |
|  | Annual Precipitation |  | Bio12 | mm | Yes |
|  | Precipitation of Wettest Month |  | Bio 13 | mm | No (High correlation with Bio 12) |
|  | Precipitation of Wettest Quarter |  | Bio 16 | mm | No (High correlation with Bio 12) |
| Vegetation | NDVI March (2017) | MODIS V6 (https://earthexplorer.usgs.gov/) | NDVI March | -1 to +1 | Yes |
|  | NDVI May (2017) |  | NDVI May | -1 to +1 | Yes |
| Demographic | Nightlight | Defence Metrological Satellite Programme-Operational Line Scan System | Nightlight | 0 to 63 | Yes |
|  | Distance from water | Waterbodies digitized in Earth Explorer and Euclidean distance calculated | Dist_Water | meter | Yes |
|  | Human Population density | SEDAC (https://sedac.ciesin.columbia.edu/) | Population Density | people/km<sup>2</sup> | Yes |
| Topographic | Elevation | ASTER DEM (https://earthexplorer.usgs.gov/) | DEM | meter | Yes |
|  | Slope |  | Slope | 0 to 90 | No (High correlation with DEM) |

Correlated variables were removed as it might lend more statistical weight to one of them, leading to model overfitting (Dormann et al., 2013). We used ‘‘Pearson correlation coefficient’’ to identify highly correlated variables, where a threshold value of 0.75 was selected to choose the final covariates (Kalboussi and Achour, 2017; Pearson et al., 2002). After initial testing of all parameters, final species distribution modeling was run using the following variables: annual mean temperature, annual precipitation, NDVI-March 2017, NDVI-May 2017, nightlight, distance from water, human population density and elevation (Table 1 and 2). Elevation was selected despite having high correlation with annual mean temperature, as it is an important ecological factor restricting swamp deer distribution within the Gangetic plains (being an obligate grassland species of the plains, swamp deer are not found in undulating, hilly areas).

**Table 2.** Multi-collinearity test by using cross-correlation (Pearson correlation co-efficient, r) among the initially selected covariates (highly correlated variables= −0.75≤Values≥0.75, highlighted in yellow) (*rejected covariates in final model).

| Layer | Bio 1 | Bio 12 | Bio 13 | Bio 16 | DEM | Dist_Water | NDVI March | NDVI May | Nightlight | Population Density | Slope |
| --- | --- | --- | --- | --- | --- | --- | --- | --- | --- | --- | --- |
| Bio 1 | 1.000 | -0.657 | -0.612 | -0.617 | -0.873 | 0.105 | -0.020 | -0.475 | -0.029 | 0.112 | -0.715 |
| Bio 12 | -0.657 | 1.000 | 0.944 | 0.983 | 0.426 | -0.100 | 0.069 | 0.415 | -0.082 | -0.120 | 0.437 |
| Bio 13* | -0.612 | 0.944 | 1.000 | 0.969 | 0.462 | -0.031 | -0.012 | 0.328 | -0.080 | -0.196 | 0.452 |
| Bio 16* | -0.617 | 0.983 | 0.969 | 1.000 | 0.437 | -0.030 | 0.028 | 0.367 | -0.078 | -0.142 | 0.443 |
| DEM | -0.873 | 0.426 | 0.462 | 0.437 | 1.000 | 0.232 | -0.143 | 0.278 | 0.035 | -0.203 | 0.774 |
| Dist_Water | 0.105 | -0.100 | -0.031 | -0.030 | 0.232 | 1.000 | -0.298 | -0.209 | -0.076 | -0.254 | 0.056 |
| NDVI March | -0.020 | 0.069 | -0.012 | 0.028 | -0.143 | -0.298 | 1.000 | 0.146 | -0.136 | 0.137 | -0.089 |
| NDVI May | -0.475 | 0.415 | 0.328 | 0.367 | 0.278 | -0.209 | 0.146 | 1.000 | 0.036 | 0.037 | 0.186 |
| Nightlight | -0.029 | -0.082 | -0.080 | -0.078 | 0.035 | -0.076 | -0.136 | 0.036 | 1.000 | 0.413 | -0.027 |
| Population Density | 0.112 | -0.120 | -0.196 | -0.142 | -0.203 | -0.254 | 0.137 | 0.037 | 0.413 | 1.000 | -0.163 |
| Slope* | -0.715 | 0.437 | 0.452 | 0.443 | 0.774 | 0.056 | -0.089 | 0.186 | -0.027 | -0.163 | 1.000 |

#### Model parameters, performance and selection

We developed and tested 10 different MaxEnt models by modifying the default parameters in order to derive the best model (Aryal et al., 2016) for swamp deer distribution. Use of default parameters can produce an over-complex or over-simplistic model (Merow et al., 2013; Warren and Seifert, 2011). MaxEnt models are sensitive to the model parameter “regularization multiplier” value (default=1), which determines the output distribution (Elith et al., 2010). A smaller parameter value will result in a more localized output distribution, while a larger value will generate a more spread out, less localized predictions (Phillips, 2017). We developed models with 10000 background sampling points as environmental space (Wilting et al., 2010) with different regularization parameter values (0.5-5). We randomly assigned the presence localities as training and testing dataset (70% as training, 30% as test). Rest of the settings for each parameter were kept default in each run. To select the best-fitted regularization multiplier, we used the Corrected Akaike information criterion (AICc) based evaluation method and selected the model with the lowest AICc value (Cordellier and Pfenninger, 2009; Williams et al., 2017). The evaluation of different regularization multiplier values has been done using the software ENMTOOLS version 1.4.4 (Warren and Seifert, 2011).

After selecting the best-fitted regularization multiplier (Model with lowest AICc value), we ran the final model with 10 replicates of 5000 iterations in MaxEnt to derive the average model (Phillips et al., 2006). An auto feature limiting function was used to fit the species environmental curves and to train the MaxEnt models. The minimum probability of MaxEnt prediction was set to “Minimum training presence Logistic threshold” (MTPLR) (Montemayor et al., 2014). We used the logistic model output which displays suitable value from 0 (unsuitable) to 1 (suitable). To check which variables were the most important for model building a Jackknife analysis was done (Phillips, 2017).

To assess qualitative characterization of the final species distribution model, we used receiver operating characteristic (ROC) area under curve (AUC) values (Phillips et al., 2006). AUC value of a model ranges from 0 to 1, where 0.90 to 1.00 range is considered as a good fit (Araújo et al., 2005; Hemsing, 2010).

#### Swamp deer validation surveys

The MaxEnt prediction gave a probability distribution map of swamp deer in the upper Gangetic plains of north India. To confirm model predictions, we conducted validation surveys in selected areas of this landscape. We selected the completely human-dominated lower part of Hastinapur Wildlife Sanctuary (downstream of Bijnor Barrage along Ganga) and Sohagibarwa Wildlife Sanctuary (in Sharda habitat block) in Uttar Pradesh for validation surveys. Hastinapur Wildlife Sanctuary is probably the most critical swamp deer habitat in the Ganga habitat block and earlier extensive surveys were conducted till Bijnor Barrage (see Paul et al., 2018). We surveyed the entire stretch of Ganga from Bijnor Barrage till the southern boundary of Hastinapur Wildlife Sanctuary (about 100 km), covering all grassland patches found on either side of the river during May 2017 and April 2018. On the other hand, Sohagibarwa Wildlife Sanctuary is the easternmost protected area predicted by the model, known to harbour large patches of grassland habitat and historical presence of swamp deer (Ghimire et al., 2019). Apart from field-based information (direct sighting, antler and genetically confirmed pellets), we have also collected secondary information (newspaper report, personal communications etc.) on swamp deer presence in the southernmost predicted regions of our study area. To assess the accuracy of model prediction, we extracted pixel values of confirmed unique (duplicates removed) swamp deer presence points (primary and secondary) and considered those points with more than Minimum training presence Logistic threshold (MTPLR). The formula used to calculate accuracy is:

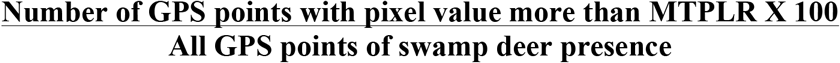

### Identification of Priority Conservation Areas

Earlier work on the *Rucervus duvaucelii duvaucelii* showed fragmented, but fine-scale distribution of the subspecies in the northern part of the Ganga habitat block (till Bijnor barrage) (Paul et al. 2018). As we generated detailed information on the subspecies’ entire range (covering both Ganga and Sharda habitat blocks) and habitat status based on the initial surveys, distribution modeling and validation surveys, it was meaningful to identify critical ‘‘Priority Conservation Areas’’ where immediate management attention is required. This entire landscape is a mosaic of protected and non-protected areas with very high pressures from anthropogenic interventions. Between these two habitat blocks there are differences in grassland coverage area, management regimes and human densities. We used a few criteria (see below) to select such ‘‘Priority Conservation Areas’’ across the entire landscape covering both protected and non-protected habitats. The criteria were: (a) presence of grassland habitats (through field surveys and GIS approaches, Paul et al. 2018); (b) results from the MaxEnt model predictions; (c) confirmed species presence in the area; and (d) strategic position of the areas within this human-dominated mosaic landscape to ensure continuation of the species’ seasonal migration (Martin and Gopal 2015; Paul et al. 2018). Most importance was given to species and grassland presence, followed by the other two criteria.

## Results

### Swamp deer initial survey in Sharda habitat block

We found much fine-scale swamp deer occurrence evidence than earlier reported (Qureshi et al., 2004) across majority of surveyed areas in the Sharda river habitat block including all the prominent protected areas (Pilibhit Tiger Reserve, Kishanpur Wildlife Sanctuary, Dudhwa National Park and Katerniaghat Wildlife Sanctuary). Additionally, we found presence of the species in non-protected grasslands of South Kheri and North Kheri Forest Divisions of Uttar Pradesh. Apart from swamp deer, we also identified the presence of other cervids (hog deer, chital, sambar etc.) in the grassland habitats. We collected a total of 225 biological samples (two swamp deer carcasses, 144 pellets and 79 confirmed swamp deer antlers) and identified 132 pellets out of which 51 belonged to swamp deer and 70, 10 and one to hog deer, chital and sambar, respectively. Remaining 12 samples did not provide any results during molecular species identification.

### MaxEnt prediction of swamp deer distribution

Out of the 10 different models with different regularization multipliers (RM 0.5-5), the model with RM 1.5 showed lowest AICc Value (1183.32) and was selected (Supplementary Table 1). We pooled the prediction probabilities (above MTPLR=0.067) to create high (0.6-1), moderate (0.4-0.6) and low probability (0.067-0.4) occurrence zones. The high probability of swamp deer was predicted to be along the banks of rivers of both Ganga and Sharda blocks (Ganga, Kotawali, Kosi, Ramganga, Sharda, Suheli, Ghaghra and Baur) (Figure 2). In the Gangetic habitat block, the highest probabilities were found to be upstream of Bijnor Barrage along river Ganga, Banganga and Solani. The moderate zones included majority of the above-mentioned areas in addition to basins of Gomti, Gaula, Devha, Baigul, Ganga River (downstream of Bijnor Barrage) and some parts of Suhelwa Wildlife Sanctuary and Sohagibarwa Wildlife Sanctuary. The low probabilities of occurrence were found to be along river Yamuna and Hindon on the west; Rapti, Kuwano and some scattered areas of Sohelwa and Sohagibarwa Wildlife Sanctuary in the east and few areas further downstream south along Ganga (Figure 2). Out of the total predicted areas for swamp deer distribution (10968 km^2^), 3413 km^2^ falls within protected areas of Uttarakhand and Uttar Pradesh.

**Figure 2:**
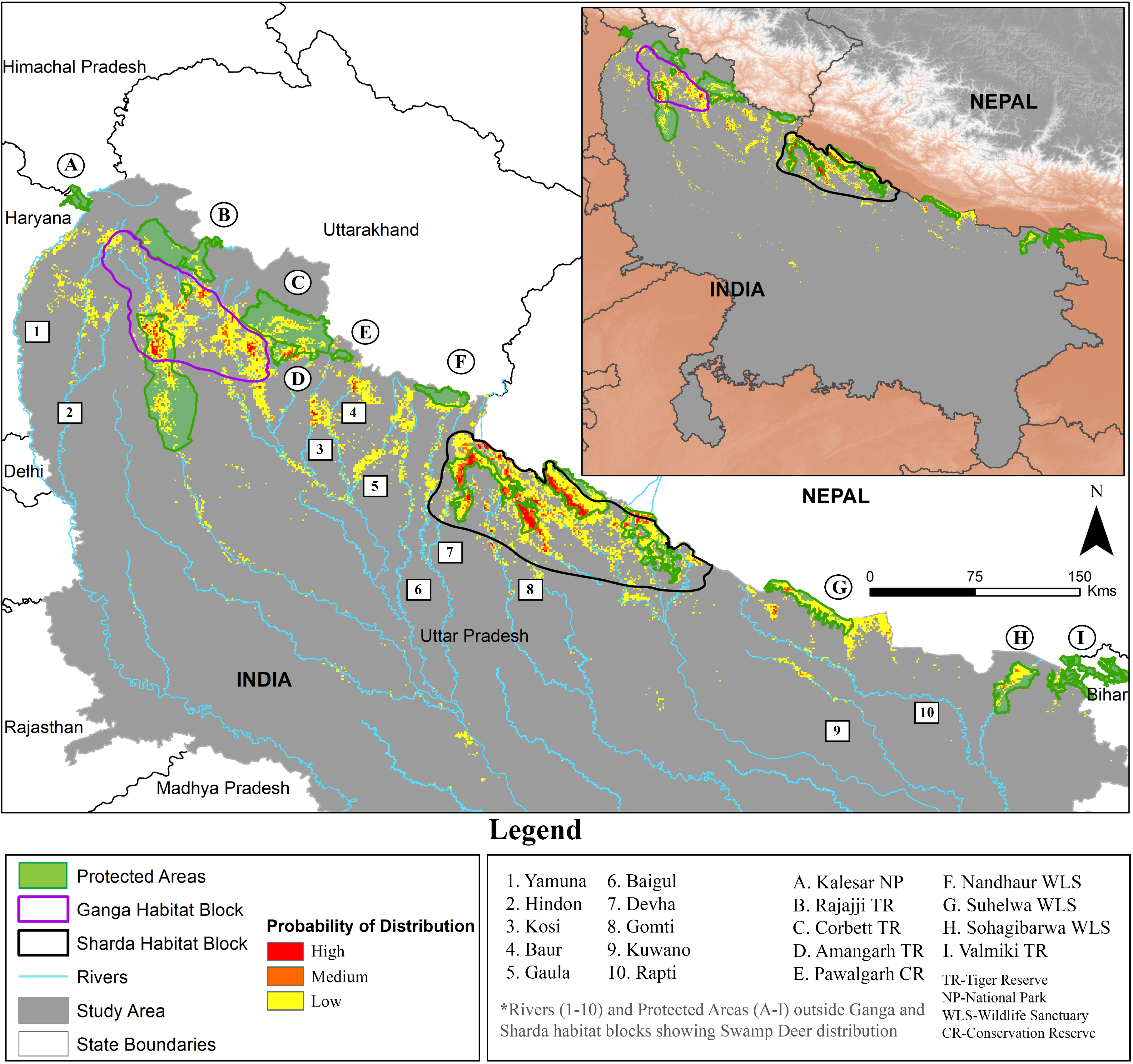
Map showing the predicted distribution of swamp deer through MaxEnt modeling. The major rivers and protected areas outside Ganga and Sharda Habitat Blocks are highlighted and marked.

The jackknife prediction of the final selected model showed that the NDVI-May had highest contribution (37.1%), followed by annual mean temperature (24.5%), distance from water (18.6%), nightlight (8%), NDVI-March 2017 (7.1%) in predicting swamp deer distribution. The remaining 4.7% is contributed by human population density, DEM and annual precipitation. Majority of the predicted areas were having an elevation of about 200m, and mean annual temperature range between 22°C-25°C. MaxEnt results showed best prediction at a NDVI value of 0.45 during March and 0.6 in case of May. Distance from water showed a negative relationship with swamp deer distribution after a distance of seven km from the rivers. Both nightlight and human population density were negatively related to the probability of distribution. The AUC value of the final selected model was 0.992 with a standard deviation of 0.001; indicating excellent prediction for swamp deer distribution (Elith et al., 2006) (Supplementary Figure 1 and 2).

### Validation surveys

Our survey in the lower part of Hastinapur Wildlife Sanctuary revealed the presence of fragmented grassland patches with swamp deer evidence (Figure 3). In addition to direct sighting (n=1) and shed antlers (n=6), molecular analysis of pellet samples (n=163) confirmed presence of swamp deer (n=82), hog deer (n=18), nilgai (n=25), barking deer (n=3) and domestic goat (n=4) (Supplementary Table 2). Rest 31 samples did not amplify during Polymerase Chain Reaction (PCR). In Sohagibarwa Wildlife Sanctuary, our surveys revealed *Typha* sp. and *Phragmites* sp. dominated grasslands but no evidence of swamp deer (Supplementary Table 2). Additionally, we validated the southernmost predicted location along Ganga (locally known as Kanauj, Figure 3) through secondary sources (direct sighting and media reports). Considering all these evidences (n=32 unique occurrence points), we obtained a model accuracy of 84.37% after ground validation. Our validation surveys approximately expand the known range of swamp deer by 1125 km^2^ along Ganga in the south.

**Figure 3:**
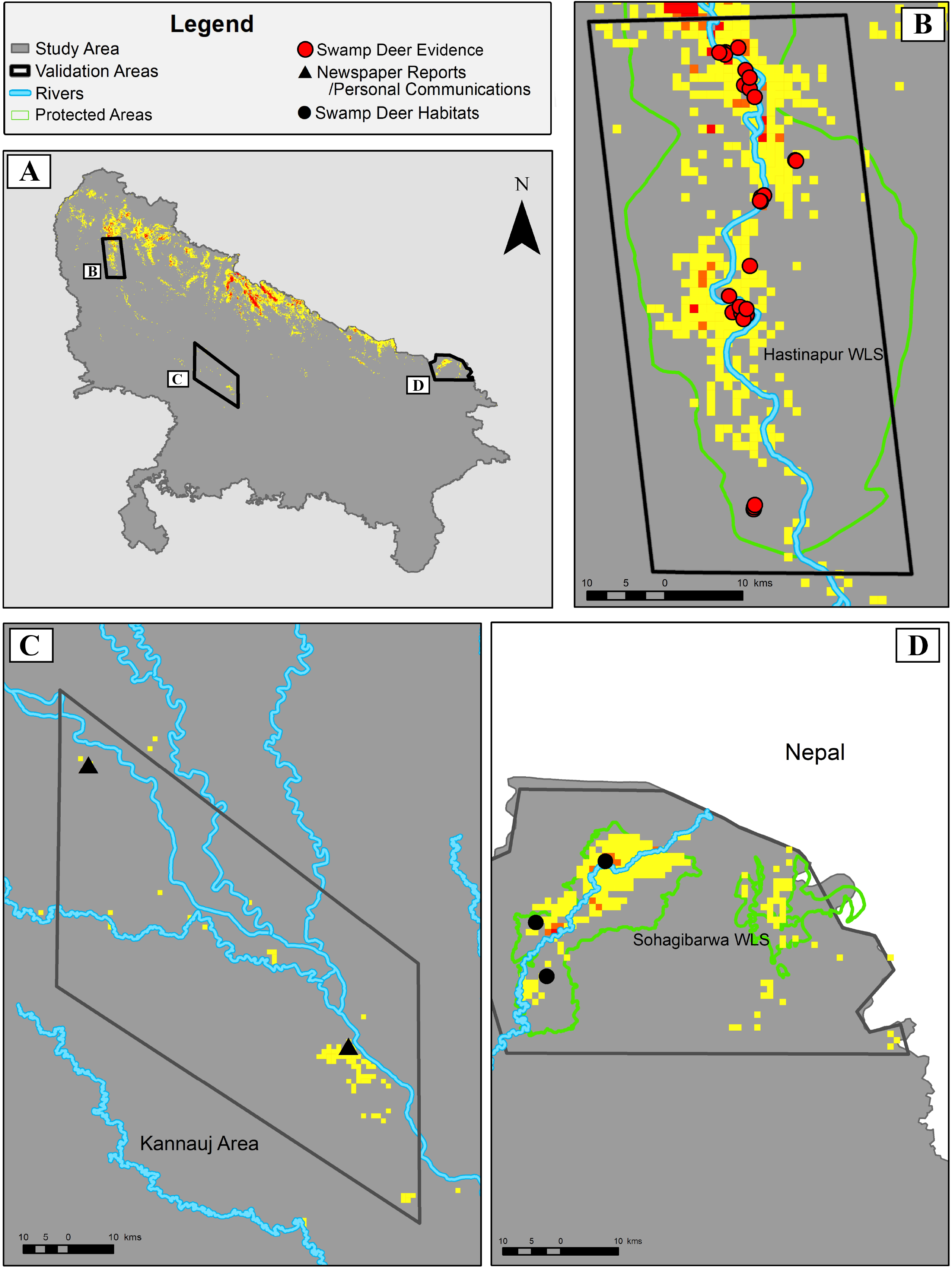
Map showing the swamp deer evidence locations within the surveyed areas predicted through MaxEnt models.Within the figure, (A) shows three selected validation areas; (B) Validation area 1-Part of Hastinapur Wildlife Sanctuary below Bijnor Barrage, Validation area 2-Kannauj along Ganga and (D) Validation area 3-Sohagibarwa Wildlife Sanctuary.

### Habitat status and disturbances

Majority of the grasslands and its fauna within the protected areas of Sharda habitat block are undisturbed and receive good protection. However, parts of Pilibhit Tiger Reserve and Katerniaghat Wildlife Sanctuary are facing anthropogenic disturbances like overgrazing and habitat conversion, and the non-protected areas in the Sharda block are experiencing rapid habitat encroachment (Supplementary Table 2). The grasslands in Sohagibarwa Wildlife Sanctuary are protected, undisturbed and hold potential for future relocation of swamp deer. Similarly, we found that the grasslands downstream of Bijnor Barrage in Hastinapur Wildlife Sanctuary (Ganga habitat block) are facing severe pressures from overgrazing, habitat conversion and occasional poaching (Supplementary Table 2). The remaining habitats found downstream river Ganga are highly fragmented and future viability of swamp deer cannot be ascertained at this point.

### Priority Conservation Areas

1. Based on the specific selection criteria (see methods section), we identified four swamp deer “Priority Conservation Areas” covering both habitat blocks in this region (Figure 4,Supplementary Table 3). Of these four selected areas, three (one protected and two non-protected areas) are in the Ganga habitat block, whereas one (non-protected areas) is in the Sharda habitat block. The details of these areas are as follows:
2. Priority Conservation Area 1: This is a ~193 km^2^ area along Ganga between Jhilmil Jheel Conservation Reserve, Uttarakhand and Hastinapur Wildlife Sanctuary, Uttar Pradesh (Figure 4A, Supplementary Table 3). We quantified ~14 km^2^ grassland habitat (~7% coverage) within this area with abundant evidence of swamp deer. The entire area is human-dominated and non-protected, and acts as a potential migration route between the above-mentioned protected areas (Paul et al. 2018).
3. Priority Conservation Area 2: This area covers the lower part of Hastinapur Wildlife Sanctuary downstream to Bijnor Barrage (1526 km^2^) (Figure 4B, Supplementary Table 3). This is a protected area with large human habitations within its boundary. We quantified ~73 km^2^ grassland habitat (~4% coverage) with swamp deer evidence from different parts.
4. Priority Conservation Area 3: This is an isolated, recently reported swamp deer population (Paul et al. 2018) along Ramganga river in Jamanpur, Afzalgarh, Uttar Pradesh (Figure 4C, Supplementary Table 3). This is a 139 km^2^ area with ~3 km^2^ grassland habitat (~2% coverage) and retains a small population of swamp deer (Paul et al. 2018).
5. Priority Conservation Area 4: This is a 555 km^2^ unprotected riverine landscape along Sharda river (Figure 4D, Supplementary Table 3). It retains ~136 km^2^ grassland habitat (~24% coverage) and connects the Shuklaphanta Wildlife Sanctuary, Nepal with Kishanpur Wildlife Sanctuary, Uttar Pradesh, and possibly acts as the main corridor to connect the Nepal population with the north Indian population.

**Figure 4:**
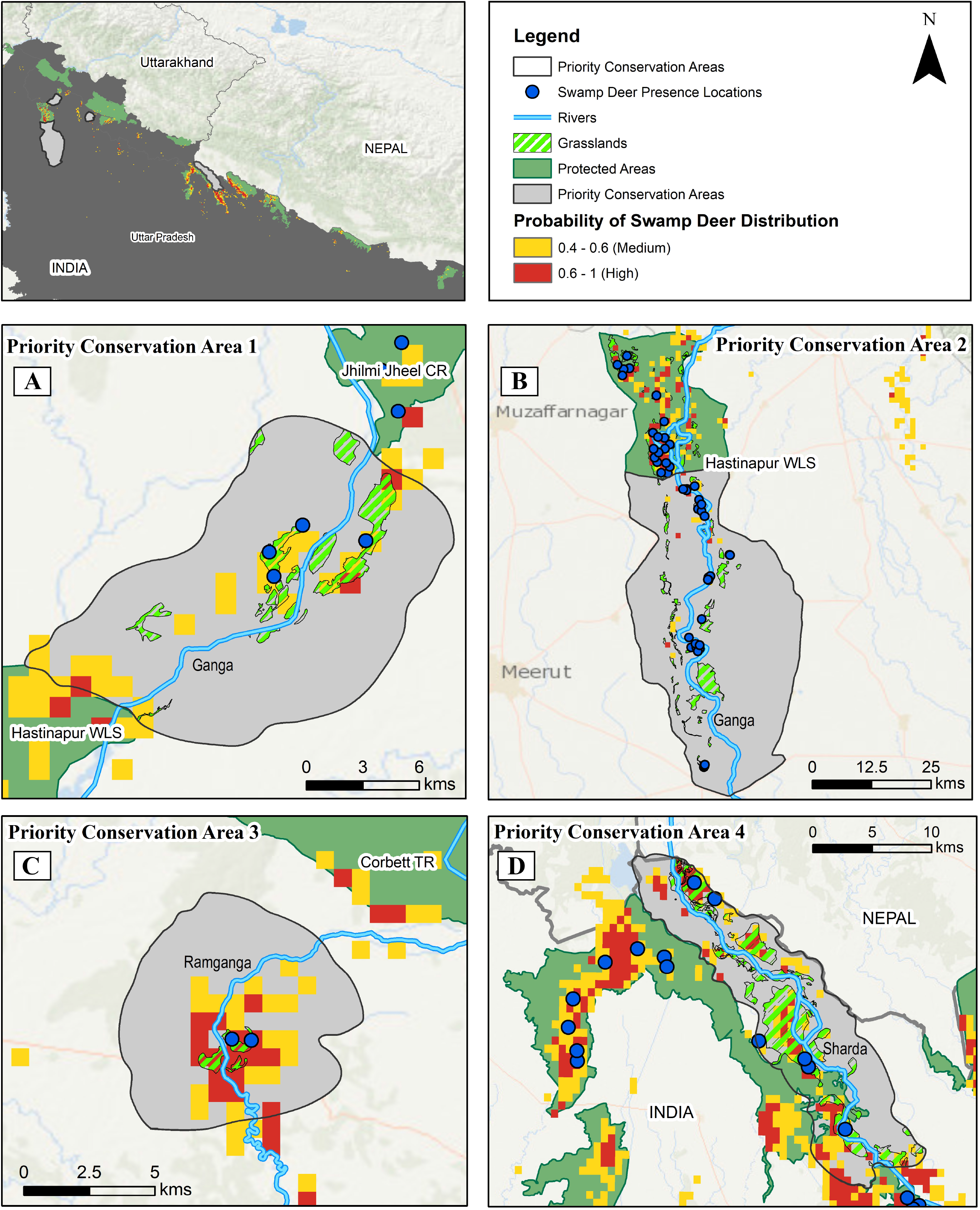
Map showing the identified four “ Priority Conservation Areas” in our study area. The priority areas identified are (A)The stretch of Ganga between Jhilmil Jheel Conservation Reserve and Hastinapur Wildlife Sanctuary; (B) The lower part of Hastinapur Wildlife Sanctuary below Bijnor Barrage; (C) The isolated grassland area on the bank of Ramganga near Jamanpur, Afzalgarh and (D) The unprotected area around Sharda river between Shuklaphanta Wildlife Sanctuary, Nepal and Kishanpur Wildlife Sanctuary, India.

## Discussion

To the best of our knowledge, this is the most exhaustive study conducted to assess the northern swamp deer (*Rucervus duvaucelii duvaucelii*) distribution using species distribution modeling and subsequent ground-truthing surveys across the subspecies’ range in India. Compared to the Ganga habitat block, our surveys show that the Sharda block still retains considerably large and undisturbed patches of grassland habitat with reports of significant swamp deer population (Duckworth et al., 2015; Qureshi et al., 2004;), thereby acting as a primary stronghold for the northern subspecies. During our surveys we found large numbers of swamp deer evidence in Pilibhit Tiger Reserve, Kishanpur Wildlife Sanctuary and Dudhwa National Park, whereas less signatures from Katerniaghat Wildlife Sanctuary. However, the non-protected areas along the rivers harbouring grasslands are experiencing a lot of disturbance and require urgent conservation attention. Anecdotal reports from some of these areas (Kishanpur Wildlife Sanctuary of India and Shukla Phanta Wildlife Sanctuary of Nepal) indicate that they are connected by regular swamp deer movements involving non-protected areas but the exact routes and patterns of habitat use are not known. Future studies should focus on using radio telemetry and genetic tools to understand these aspects. The Sharda habitat block is an integral part of contiguous central Terai landscape and hosts the majority of the swamp deer population (Qureshi et al., 2004; Singh, 1978). It will be important to focus on estimating population size, assess critical viable habitats and corridors in this landscape for future conservation of swamp deer.

During this study, we have carefully designed the swamp deer distribution modeling and used relevant parameters based on the species ecology. Although it is known that species distribution modeling for habitat-specialists yields better predictions (Connor et al., 2017; Rhoden et al., 2017), it is critical to have a focal area-based modeling approach, selection of ecologically meaningful covariates and accurate presence locations (Araújo and Guisan, 2006; Costa et al., 2015; Tye et al., 2017; Williams et al., 2012). We used only confirmed primary data (direct sightings, antlers and genetically identified pellets) in the models to reduce false predictions due to sample biases resulting from inaccurate information (historical records, personal communications etc.) (Tye et al., 2017). We ensured accurate species identification for every presence location as earlier studies suggested erroneous species identification can have severe downstream impacts in data interpretations (Aubry et al., 2017; Beerkircher et al., 2009; Costa et al., 2015). Further, we conducted the entire analysis at a spatial scale of one km^2^ (corresponding to average grassland patch size in this landscape) to avoid model overfitting (Tye et al., 2017) and used most recent covariate information to maintain temporal correspondence between the occurrence points and the variables (Phillips et al., 2006). It is important to point out that we did not use species absence data during analyses as it was challenging to determine true absence from these mosaic habitats consisting of grassland and croplands. However, we think that for swamp deer this would not have any impact on the results as the species is only restricted within 5-10 km on either side of the rivers present in this region (Paul et al., 2018; this study). Overall, our model showed very accurate predictions (AUC value of 0.992), which is expected for a habitat-specialist species (Connor et al., 2017) (Supplementary Figure 2). Finally, we conducted extensive validation surveys to confirm the accuracy of the distribution predictions by the models. Although most of the species distribution modeling studies result in identification of new areas of occurrences (Hernandez et al., 2006), the validations are largely restricted to comparison of training data subsets with predictions (Fois et al., 2018; Pearson et al., 2007) and not on-field validations (Greaves et al., 2006; Jiménez-Valverde et al., 2011). Considering all these points, we feel that our approach has minimized biases in distribution predictions arising from various factors (for example small sample size- Yañez-Arenas et al., 2014; biased sampling- Syfert et al., 2013; lack of essential variables- Barry and Elith, 2006; species misidentification- Costa et al., 2015; ambiguous locations- Moudrý and Šímová, 2012; lack of on-field validations- Mccarthy et al., 2012 etc.).

Out of the 11 initial ecological/environmental covariates (eight in the final model) used, NDVI May, annual mean temperature, distance from water, nightlight and NDVI March were found to be the most important factors (~95%) in explaining swamp deer distribution (Supplementary Figure 1 and 2). In the entire Gangetic plains, the croplands possess a lionshare of the Landuse Landcover in terms of area. Thus we chose NDVI May and NDVI March which are extremely critical habitat covariates that represent the grasslands-cropland mosaics, where the March NDVI indicates indices of both vegetation types during the pre-harvesting period when it is difficult to differentiate them (thus overlapping NDVI values). However during May, after harvestation of the crops (wheat and sugarcane) the same NDVI values are only represented by grasslands. As swamp deer is an obligatory grassland species, it makes perfect ecological sense that NDVI May showed strong positive relation with probability of swamp deer presence and was the highest contributor in explaining the species distribution. We used NDVI as a surrogate for grassland delineation in our analyses (comparatively lower accuracy) as detailed ground-truthed visual classification of the grasslands was not available. In the absence of true grassland covariate, we interestingly found low prediction probabilities in Suhelwa Wildlife Sanctuary though previous reports suggest absence of tall wet swampy grasslands (Chanchani et al., 2014). The prediction has been driven majorly by distance to rivers and NDVI as the area has a lot of rivers flowing through the sanctuary but further surveys should be conducted to ascertain the current status. Thus, future studies should focus on generating fine-scale ground validated information on grasslands for further fine-tuning of the models. However, it is important to remember that the Gangetic basin is an extremely dynamic landscape and locations of these small grassland patches can change over a short span of time. Surveys at regular intervals to identify and mapping these habitats will be very critical.

Our validation surveys confirmed swamp deer presence in the lower part of Hastinapur Wildlife Sanctuary and identified only grassland habitats in Sohagibarwa Wildlife Sanctuary. During the surveys, we identified highly fragmented habitat patches with comparatively less swamp deer evidence below the Bijnor Barrage area of Hastinapur Wildlife Sanctuary along river Ganga. This area is highly human-dominated and needs immediate conservation attention for future survival of the species. There are sporadic reports of swamp deer presence further south along Ganga (Figure 3) and extensive surveys need to be conducted to ascertain viable habitat and population status. Within Sohagibarwa Wildlife Sanctuary we identified large patches of wet grasslands without any swamp deer evidence. Earlier studies reported swamp deer presence in this region during the 1960s (Ghimire et al., 2019) and given the protection status of this sanctuary, it can be a suitable area for potential reintroduction in future.

Finally, we identified four critical “Priority Conservation Areas” that demand immediate management attention for long-term survival of the species in this landscape (Figure 4). All of these selected areas are either corridor habitat (Priority Conservation Area 1 and 4, Figure 4A and 4D) or important populations that need careful management. Three of these habitats are in the Ganga habitat block, where habitat loss and other anthropogenic factors have resulted in extremely fragmented swamp deer habitats in recent time (Paul et al. 2018). The Priority Conservation Area 1 (between Jhilmil Jheel Conservation Reserve and Hastinapur Wildlife Sanctuary, Figure 4A) is the only connecting habitat between these protected regions. Despite having unprotected status, this area has a grassland habitat coverage of ~7% which is similar to Jhilmil Jheel Conservation Reserve and Hastinapur Wildlife Sanctuary (3% and 7%, respectively) and retains swamp deer evidence. Swamp deer from Jhilmil Jheel Conservation Reserve are known to migrate seasonally (Paul et al. 2018) and most likely use this area to move downstream Ganga. However, this area currently faces a lot of human disturbances (Paul et al. 2018) and if these grassland habitats are lost, the two protected areas will become isolated. Conservation efforts should strongly focus on public awareness programs and community involvement through developing this area as a ‘‘Conservation or Community Reserve’’ (based on existing land use and land-tenure data) in near future to ensure long-term survival of this subspecies. However, in case of the Priority Conservation Area 2 (downstream to Bijnor barrage area, Figure 4B) the challenges are different. This region is within a protected area (Hastinapur Wildlife Sanctuary) and thus the species status should be better here. A large part of the sanctuary (particularly downstream of Bijnor Barrage) is heavily human-dominated (including large villages and towns) and possibly cannot be reverted back to natural grassland habitats. We suggest further detailed, fine-scale mapping of this region to demarcate the human-dominated areas and rationalize the boundary to include only grassland habitats for focused habitat management. The third Priority Conservation Area is an isolated swamp deer population in the eastern border of the Ganga habitat block (Figure 4C). This population is completely surrounded by human habitats and needs urgent attention. This area is also the closest to the Sharda habitat block populations and future investigation should focus on understanding the historical connectivity between these two habitat blocks. Within the Sharda habitat block we identified one Priority area that is a critical migration corridor to connect the Nepal population with the Indian swamp deer population. While most of the swamp deer habitat in this block is within a number of protected areas and thus receive better protection (unlike the Ganga habitat block), this critical corridor is mostly unprotected. Given that this is the world’s largest global intermixing population of swamp deer, protecting this corridor by converting this area as a ‘‘Conservation or Community Reserve’’ possibly will be the only way to maintain this metapopulation. If this corridor is lost due to human impacts, the Indian population might get genetically isolated from the Nepal population.

It is noteworthy to point out that the importance of these priority conservation areas should not be misconstrued as purely based on grassland coverage/extent alone. The fragmented grassland habitats outside protected areas are easy targets for habitat conversion in this high human-dense landscape, and majority of the selected priority areas have low overall grassland coverages (~2-24% of the respective areas, Supplementary Table 3). This can be argued against our suggestion of creating priority conservation areas. However, these areas have critical importance when looked at swamp deer specific requirements (essential habitat for seasonal migration, feeding ground, fawning etc.) and they are essential to ensure long-term persistence of the species in this landscape. Thus, they need to be considered as critical parts of the entire landscape for swamp deer survival. According to recent assessments, grasslands are one of the most marginalised ecosystems in India (Task Force on Grasslands and Deserts report, 2006), with a loss of ~20 million hectare area between 1880-2010. With the ‘‘Green Revolution’’ movement in India, there has been a major shift to irrigation based agricultural practises, leading to fast decline in grassland habitats (Vanak et al., 2017) across north India. These remaining grassland patches are the last resort to the habitat-specialist biodiversity, and appropriate management interventions are required urgently to protect the ecosystem.

## Conclusion

During the early 20th century, the swamp deer was found throughout the Indo-Gangetic plains (Duckworth et al., 2015) but they are currently considered to have the highest extinction probability among all Indian megaherbivores (Karanth et al., 2010). However, details of their population size and habitat status have been rather scanty due to logistical challenges in generating population level information in human-dominated landscapes across the Gangetic plains. The result of our study provides a detailed insight of swamp deer distribution and highlights the importance of the riverine-grassland systems of northern India, where a considerable amount of biodiversity co-exists with a very high human population (Uttar Pradesh ranks first among the states of India according to Census 2011). While all these new information is quite promising for the species, there is still a serious need to focus on population enumeration and grassland management, particularly those present in the priority conservation areas described here. Such efforts would also generate critical information for other sympatric species (for example, hog deer, fishing cat, Bengal florican etc.) of the grasslands/wetlands and help in their conservation. We suggest using swamp deer as an umbrella species to strengthen habitat management and conservation for the otherwise marginalized grassland habitats in this landscape. This work also emphasizes the need for similar studies to generate critical information for other locally or regionally endemic, habitat-specialist species that occur outside protected areas for their conservation.

## Acknowledgement

We acknowledge the Forest Departments of Uttarakhand and Uttar Pradesh for providing necessary permits to carry out the research. Our thanks to the Forest Department officials and local community members for their assistance during the surveys. We acknowledge help from Suvankar, Imam, Ranju, Bhura, Annu, Ammi and Prajok for their help during field surveys. We appreciate technical help from Laxminarayan, Sohini and Tathagata (GIS work) and Madhanraj, Garima and Mouli (laboratory work). We thank the Director, Dean and the Wildlife Forensics and Conservation Genetics Cell of Wildlife Institute of India for their support.

The work was funded by Uttarakhand Forest Department, Uttar Pradesh Forest Department and Ministry of Environment, Forests and Climate Change, Government of India (Grant No. 244/2018/RE). Shrutarshi Paul was awarded Department of Science and Technology INSPIRE Research Fellowship (IF150680) and Samrat Mondol was supported by Department of Science and Technology INSPIRE Faculty Award (IFA12-LSBM-47).

---

**Supplementary Figure 1:**
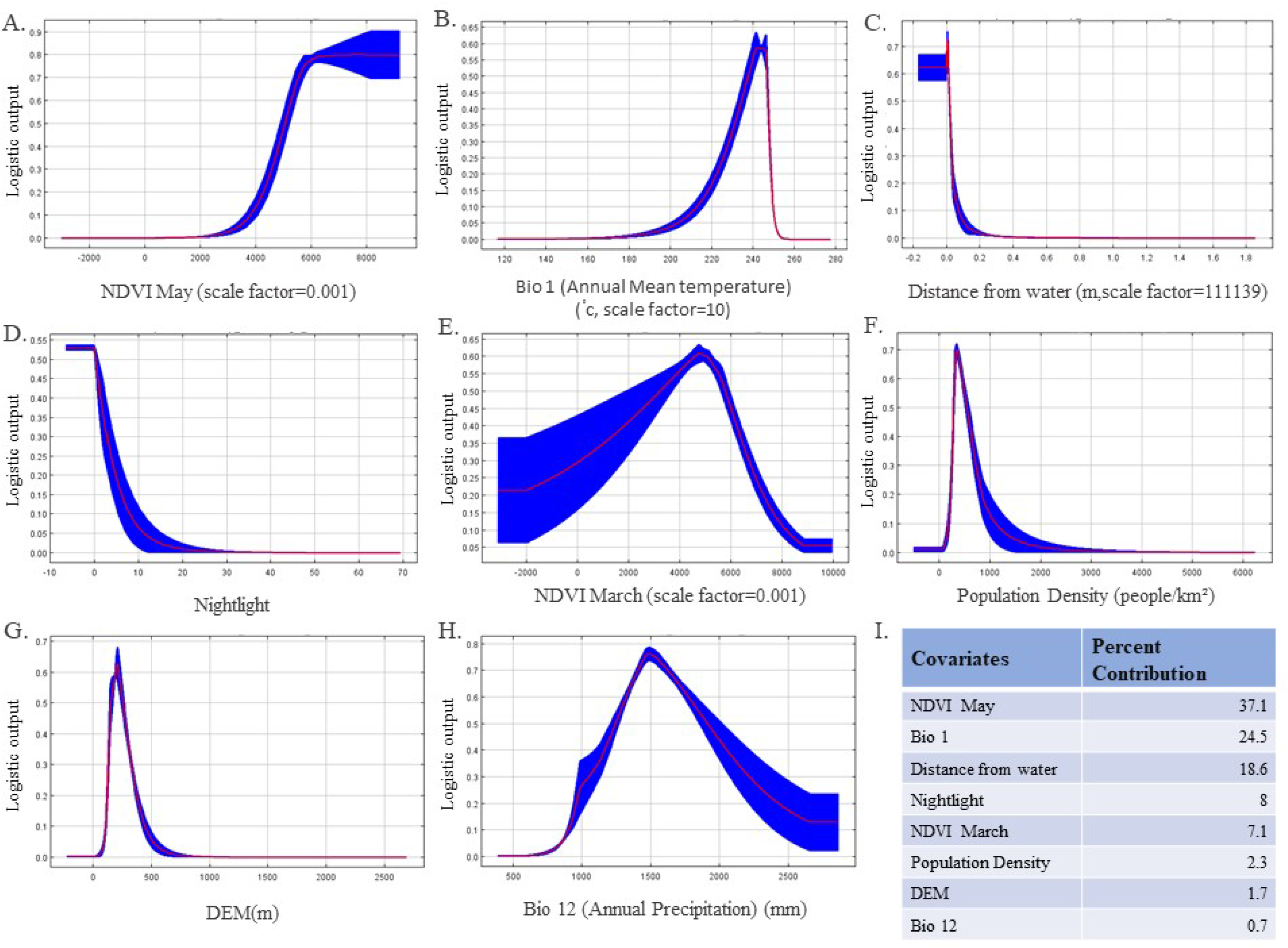
Figure showing the response curves of eight covariates used in the final MaxEnt model for swamp deer distribution predictions. Figures A-H shows the response curves, whereas I lists the estimates of relative contributions of the respective covariates in the MaxEnt predictions.

**Supplementary Figure 2:**
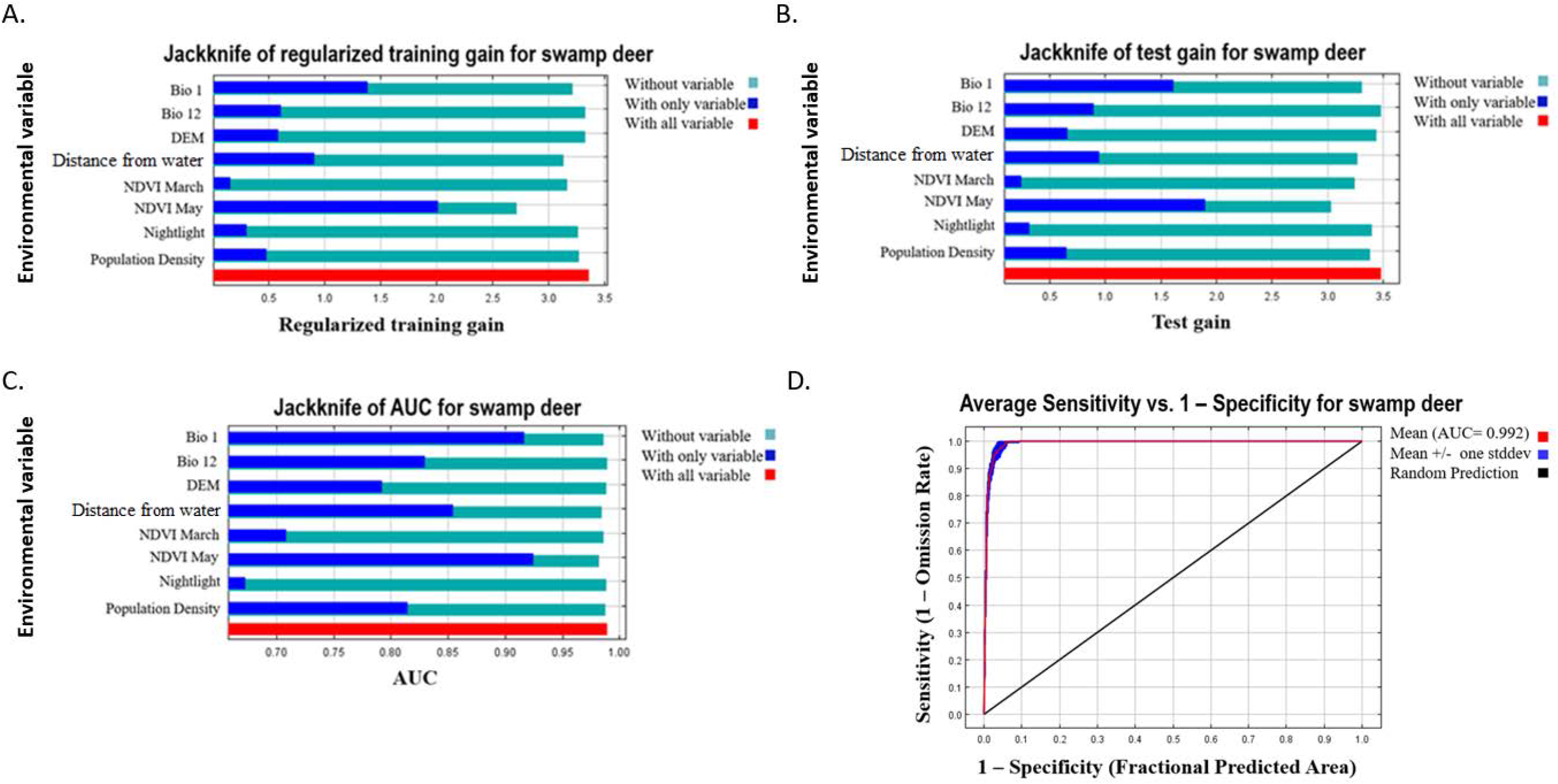
Jackknife tests of variable importance and Receiver operating characteristic (ROC) curve for predicted distribution of swamp deer, (A) Jackknife test using training gain, (B) Jackknife test using test gain, (C) Jackknife test using AUC on test data, (D) ROC curve for the training omission rate and predicted area averaged over the replicate runs.

**Supplementary Table 1:**
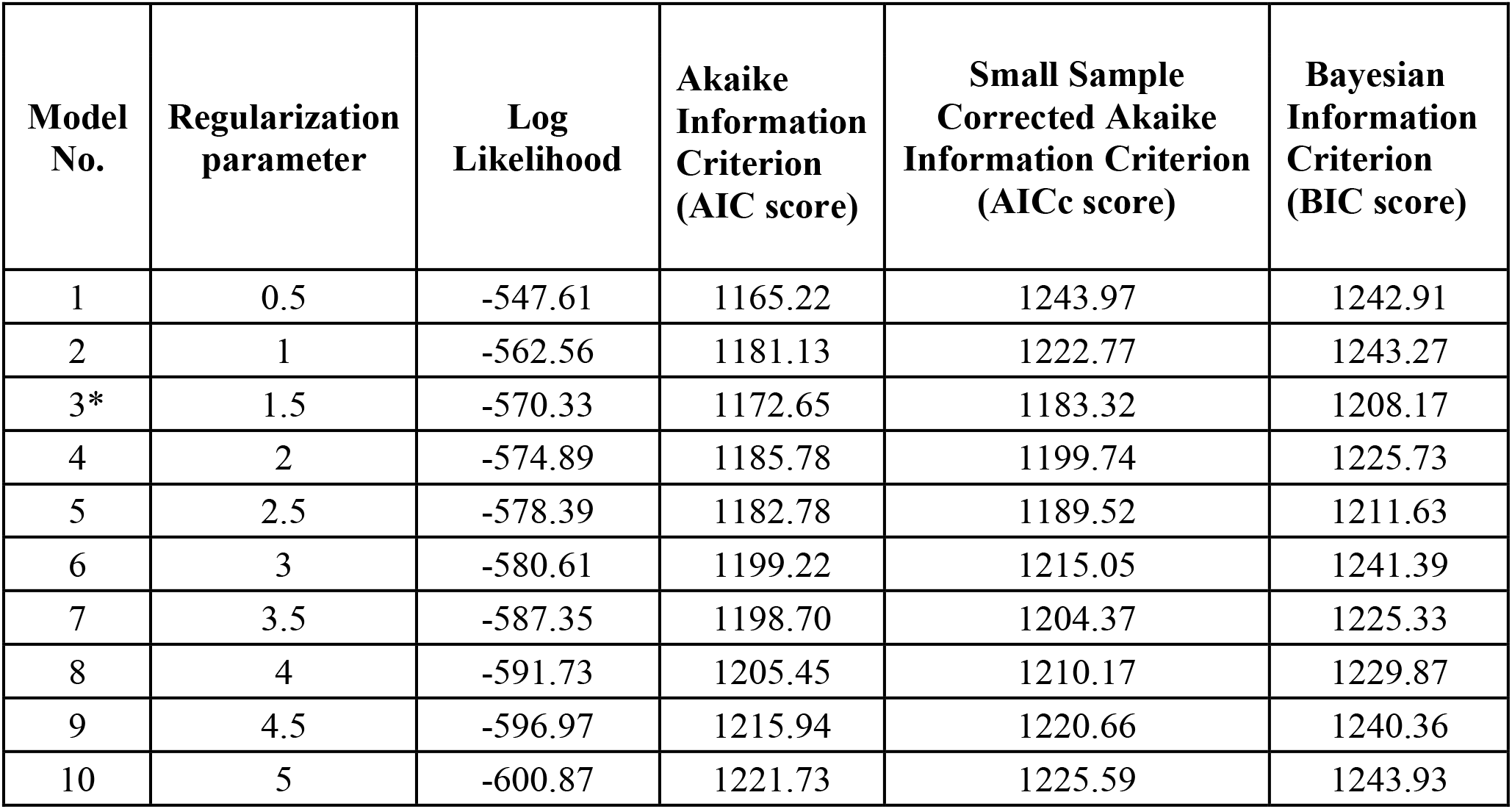
Model selection based on lowest Small Sample Corrected Akaike Information Criterion (AICc) value (*selected model)

| **Model No.** | **Regularization parameter** | **Log Likelihood** | **Akaike Information Criterion (AIC score)** | **Small Sample Corrected Akaike Information Criterion (AICc score)** | **Bayesian Information Criterion (BIC score)** |
| --- | --- | --- | --- | --- | --- |
| 1 | 0.5 | -547.61 | 1165.22 | 1243.97 | 1242.91 |
| 2 | 1 | -562.56 | 1181.13 | 1222.77 | 1243.27 |
| 3* | 1.5 | -570.33 | 1172.65 | 1183.32 | 1208.17 |
| 4 | 2 | -574.89 | 1185.78 | 1199.74 | 1225.73 |
| 5 | 2.5 | -578.39 | 1182.78 | 1189.52 | 1211.63 |
| 6 | 3 | -580.61 | 1199.22 | 1215.05 | 1241.39 |
| 7 | 3.5 | -587.35 | 1198.70 | 1204.37 | 1225.33 |
| 8 | 4 | -591.73 | 1205.45 | 1210.17 | 1229.87 |
| 9 | 4.5 | -596.97 | 1215.94 | 1220.66 | 1240.36 |
| 10 | 5 | -600.87 | 1221.73 | 1225.59 | 1243.93 |

**Supplementary Table 2:**
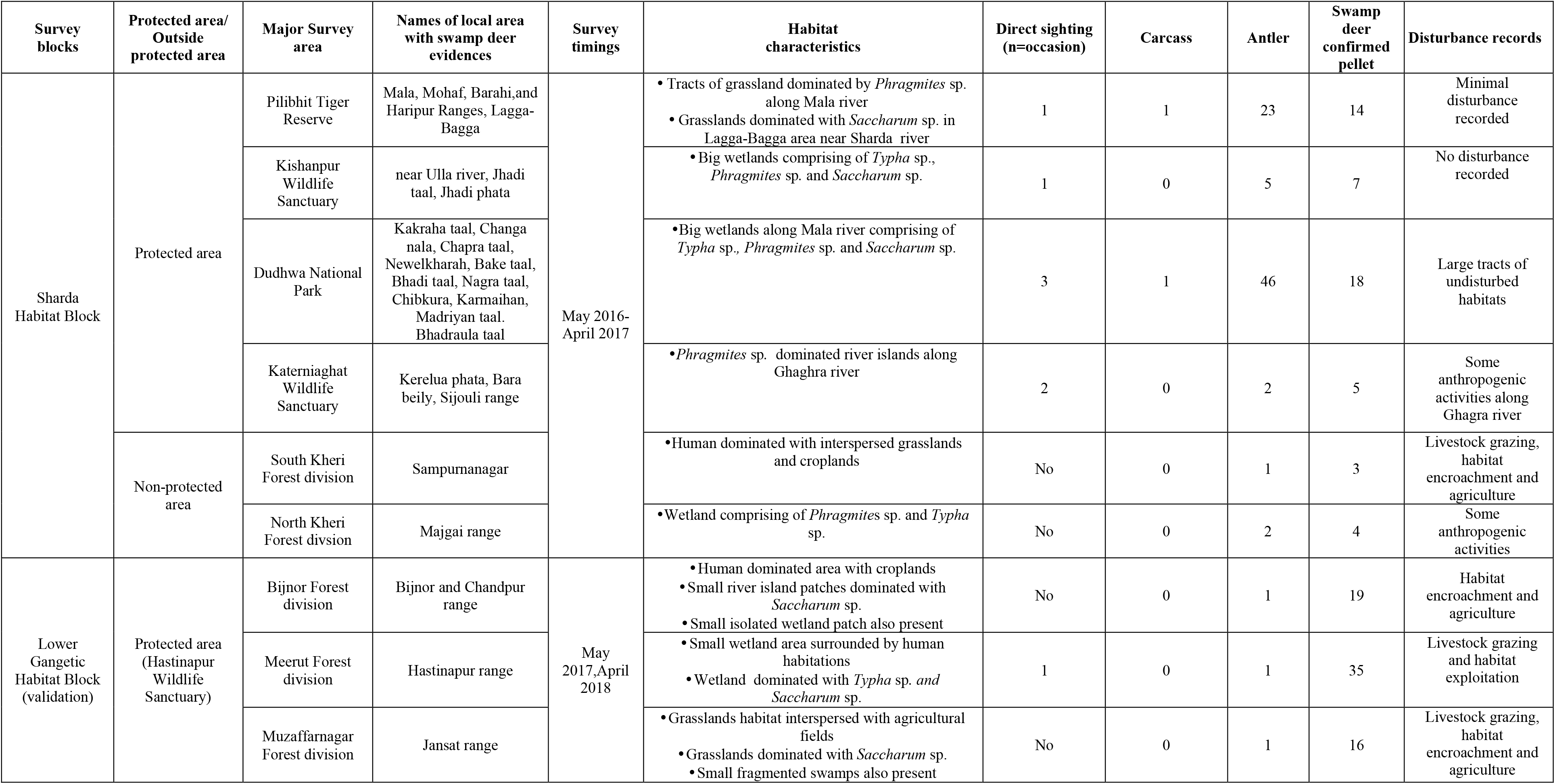
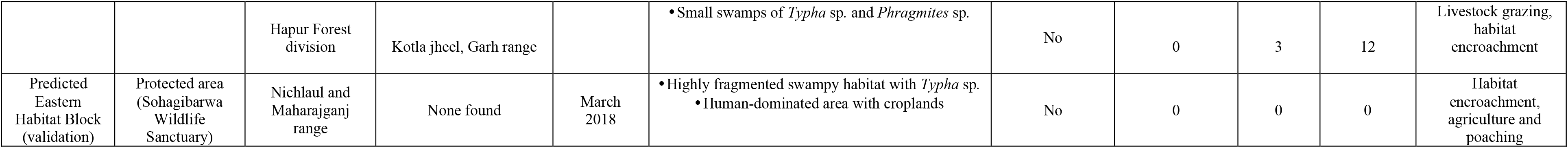
Details of swamp deer surveys in the Sharda, Lower Gangetic block and Sohagibarwa Wildlife Sanctuary using May 2016-April 2018. Information on block-wise area, habitat characteristics, direct and indirect evidences of swamp deer presence and disturbance records are presented for each survey block.

| Survey blocks | Protected area/<br>Outside protected area | Major Survey area | Names of local area with swamp deer evidences | Survey timings | Habitat characteristics | Direct sighting (n=occasion) | Carcass | Antler | Swamp deer confirmed pellet | Disturbance records |
| --- | --- | --- | --- | --- | --- | --- | --- | --- | --- | --- |
| Sharda Habitat Block | Protected area | Pilibhit Tiger Reserve | Mala, Mohaf, Barahi, and Haripur Ranges, Lagga-Bagga | May 2016-April 2017 | • Tracts of grassland dominated by *Phragmites* sp. along Mala river<br>• Grasslands dominated with *Saccharum* sp. in Lagga-Bagga area near Sharda river | 1 | 1 | 23 | 14 | Minimal disturbance recorded |
|  |  | Kishanpur Wildlife Sanctuary | near Ulla river, Jhadi taal, Jhadi phata |  | • Big wetlands comprising of *Typha* sp., *Phragmites* sp. and *Saccharum* sp. | 1 | 0 | 5 | 7 | No disturbance recorded |
|  |  | Dudhwa National Park | Kakraha taal, Changa nala, Chapra taal, Newelkharah, Bake taal, Bhadi taal, Nagra taal, Chibkura, Karmaihan, Madriyan taal. Bhadraula taal |  | • Big wetlands along Mala river comprising of *Typha* sp., *Phragmites* sp. and *Saccharum* sp. | 3 | 1 | 46 | 18 | Large tracts of undisturbed habitats |
|  |  | Katerniaghat Wildlife Sanctuary | Kerelua phata, Bara beily, Sijouli range |  | • *Phragmites* sp. dominated river islands along Ghaghra river | 2 | 0 | 2 | 5 | Some anthropogenic activities along Ghagra river |
|  | Non-protected area | South Kheri Forest division | Sampurnanagar |  | • Human dominated with interspersed grasslands and croplands | No | 0 | 1 | 3 | Livestock grazing, habitat encroachment and agriculture |
|  |  | North Kheri Forest division | Majgai range |  | • Wetland comprising of *Phragmites* sp. and *Typha* sp. | No | 0 | 2 | 4 | Some anthropogenic activities |
| Lower Gangetic Habitat Block (validation) | Protected area (Hastinapur Wildlife Sanctuary) | Bijnor Forest division | Bijnor and Chandpur range | May 2017, April 2018 | • Human dominated area with croplands<br>• Small river island patches dominated with *Saccharum* sp.<br>• Small isolated wetland patch also present | No | 0 | 1 | 19 | Habitat encroachment and agriculture |
|  |  | Meerut Forest division | Hastinapur range |  | • Small wetland area surrounded by human habitations<br>• Wetland dominated with *Typha* sp. and *Saccharum* sp. | 1 | 0 | 1 | 35 | Livestock grazing and habitat exploitation |
|  |  | Muzaffarnagar Forest division | Jansat range |  | • Grasslands habitat interspersed with agricultural fields<br>• Grasslands dominated with *Saccharum* sp.<br>• Small fragmented swamps also present | No | 0 | 1 | 16 | Livestock grazing, habitat encroachment and agriculture |
|  |  | Hapur Forest division | Kotla jheel, Garh range |  | • Small swamps of *Typha* sp. and *Phragmites* sp. | No | 0 | 3 | 12 | Livestock grazing, habitat encroachment |
| Predicted Eastern Habitat Block (validation) | Protected area (Sohagibarwa Wildlife Sanctuary) | Nichlaul and Maharajganj range | None found | March 2018 | • Highly fragmented swampy habitat with *Typha* sp.<br>• Human-dominated area with croplands | No | 0 | 0 | 0 | Habitat encroachment, agriculture and poaching |

**Supplementary Table 3:** Attributes of selection criteria for identified “Conservation Priority Areas”

| Priority Conservation Area | PCA area (km<sup>2</sup>) | Number of swamp deer evidences | Grassland coverage | Area covered by high and moderate prediction values | Strategic location importance |
| --- | --- | --- | --- | --- | --- |
| Area 1 | 193 | 22 | 7% | 13% | Connecting Hastinapur WLS and Jhilmil Jheel CR |
| Area 2 | 1526 | 89 | 4% | 2% | Important protected area of Ganga block |
| Area 3 | 139 | 20 | 2% | 26% | Isolated patch |
| Area 4 | 555 | 18 | 24% | 26% | Connecting Kishanpur WLS in India to Shuklaphanta WLS in Nepal |

## Notes

### Competing Interest Statement

The authors have declared no competing interest.

